# Small Extra-Large GTPase-like proteins influence rhizobial symbiosis in *Lotus japonicus*

**DOI:** 10.1101/2025.06.18.660248

**Authors:** Lorenzo J. Washington, Tomo Yoshino, Thalissa Malagoli Franzon, Victoria Vera, Henrik V. Scheller

**Affiliations:** Joint BioEnergy Institute, Lawrence Berkeley National Laboratory, Berkeley, California, USA; University of California-Berkeley, Department of Plant & Microbial Biology, Berkeley, California, USA; University of California-Berkeley, Department of Molecular & Cell Biology, Berkeley, California, USA

**Keywords:** GTPase, G-protein, nodulation, plant-microbe interactions, cell signaling

## Abstract

Plants possess a unique class of heterotrimeric Gα subunits called extra-large GTPases (XLGs) which contribute to numerous developmental and stress responses. In addition to the canonical Gα domain, XLGs have an uncharacterized N-terminal domain and functions that are distinct from conventional Gα subunits. In this study, we identified homologs of XLG3 in *Lotus japonicus* responsive to rhizobial and mycorrhizal symbiosis. However, these proteins were approximately one-third the size of conventional XLGs and only aligned to the N-terminal domain, containing a putative nuclear localization signal and a cysteine-rich domain of unknown function. Multiple sequence alignment and phylogenetic analysis determined these small XLGs (SXLGs) did not share domains with other mono– or heterotrimeric G-protein classes and exhibited a pattern of duplication and neofunctionalization typical of genes involved in symbiotic signaling pathways. Transient expression of *LjSXLG*s in tobacco demonstrated their potential for localization to the plasma membrane, nucleus, and nucleolus. Analysis of *L. japonicus sxlg2* mutants revealed transient impairment of immature nodule formation in a destructive experimental setup and inhibition of infection events in a nutrient-limited non-destructive experimental setup, with a delayed onset of established infection events and a potential impact on nodule maturation rate. Additionally, *sxlg2* mutants showed a potential impairment of the root growth response in N-limited conditions. We discuss the potential utility of SXLGs in better understanding the evolution of XLGs and their possible function as transcriptional regulators, as well as the likelihood SXLGs are involved in the establishment of rhizobial and mycorrhizal symbioses through influencing membrane reorganization, such as during infection thread development.

## Introduction

Plants possess heterotrimeric guanosine triphosphate (GTP)-binding proteins that have roles in relaying extracellular signals into their downstream pathways.(1) Similar to animal and fungal systems, this heterotrimeric protein is composed of Gα, Gβ, and Gγ subunits which disassociate upon activation to relay signal transduction; however plants possess distinct elements such as alternative regulation strategies, unique Gγ subunit types, and an additional class of Gα subunits in Extra-Large Gα-proteins (XLGs).(1,2) XLGs are composed of an uncharacterized N-terminal domain, possessing a nuclear localization signal (NLS) and a cysteine-rich domain of unknown function (DUF), and a C-terminal domain matching conventional Gα subunits.(2,3) The XLGs have been demonstrated to provide the diversity that was previously thought to be lacking in plant Gα subunits, with distinct and overlapping roles and functions compared to conventional Gα subunits.(1,3,4)(1,3,4)(1,3,4)

G-proteins have been found to be involved in a wide variety of plant signaling pathways and utilize the diversity present amongst subunit classes to effectively respond across such a range of signals.(3–7) Interestingly, despite operating on a timescale of seconds to minutes, plant G-proteins have been primarily associated with slow-scale developmental responses, such as organ development and cell elongation.(8,9) However, they are also implicated to participate in responses to biotic interactions, such as in immunity and symbiotic association with nodule-inducing rhizobia, processes which involve rapid intracellular responses to external stimuli.(3,5,6,10) Additionally, XLGs were shown to be involved in numerous pathways that incorporate environmental stresses, such as nutrient deficiency and infection, with developmental outcomes in above and below-ground tissues.(3,4,11)

This places XLGs as a potentially fruitful target of investigation regarding symbiotic associations in plant root systems, as these require the incorporation of numerous (a)biotic signals to determine large-scale developmental changes.(12,13) In this study, we used LotusBase (14,15) to identify two homologs of XLG3 in *Lotus japonicus* which exhibited expression patterns specific to stages of rhizobial and mycorrhizal symbiosis. However, the identified proteins — further referred to as small XLGs (SXLGs) — were approximately one-third the size of conventional XLGs and aligned to the N-terminal domain, possessing putative NLS and the cysteine-rich DUF. Multiple sequence alignment and phylogenetic analysis determined SXLGs did not share domains with similarly sized G-protein classes, such as Gβ and Gγ subunits or small monomeric GTPases such as ROP, and exhibited a pattern of duplication and neofunctionalization typical of genes involved in symbiotic signaling pathways. Transient expression of SXLGs demonstrated their potential for localization to the plasma membrane, nucleus, and nucleolus. Analysis of *Lotus japonicus sxlg2* mutants revealed transient impairment of immature nodule formation in a destructive experimental setup. Further experiments in a nutrient-limited, non-destructive setup demonstrated *sxlg2* mutants experienced a delay in and reduction of established infection events as well as a potential inhibition of nodule maturation rate. Additionally, *sxlg2* mutants showed a potential impairment of the root growth response in N-limited conditions.

## Materials & Methods

### Identification of candidate genes in LotusBase, Multiple Sequence Alignment, and Phylogenetic Tree Generation

LotusBase was used to BLAST the peptide sequence for *Lj*XLG3.(14,15) The top ten homologs were checked for transcriptional expression responses to symbiotic conditions using the ExpressionAtlas (ExPat) in LotusBase. Peptide sequences for *LjSXLG1* (LotjaGi1g1v0106400) and *LjSXLG2* (LotjaGi6g1v0043000) were aligned with Mega-X software: Muscle alignment and tree generation using UPGMA cluster methods.(16–19) The additional sequences were collected via pBLAST using the non-redundant protein database in NCBI.(20)

### Germination and Growing Conditions of *Lotus japonicus*

Seeds were scarified in ≥98% concentrated sulfuric acid for 25 min at 28°C and 600 rpm before 5 washes with deionized (DI) water. Seeds were surface sterilized in 10% bleach v/v for 2 minutes while shaking by hand before another 5 DI water washes. A final volume of DI water was added and seeds were left to imbibe while rotating at room temperature for 2-4 h. Afterwards, they were placed on 1/2 Murashige and Skoog (MS), 1% w/v plant tissue culture (PhytoTech Labs A111) agar plates with moistened filter paper to maintain humidity. The plates were sealed with parafilm and placed in a growth chamber (Percival Scientific: AR-100L3) set to 16:8h light:dark regime (Hi Point Z4 Control LED Sunlight), 22°C, and 60% relative humidity for 7 days before transplanting. All experiments using *Lotus japonicus* were done in these growing conditions. Wildtype Gifu and *Ljsxlg2* (LORE1 ID: 30063804; LotjaGi6g1v0043000) seeds were given the same treatment.

### Identification of homozygous *sxlg2* mutants

Wildtype Gifu and LORE1 transposon mutagenized seeds were obtained from LotusBase. The LORE1 line (30063804) R3 generation seeds of transposon mutagenized *L. japonicus* were germinated and grown for 7-14 days as described.(21) Homozygous transposon insertions in *SXLG2* were confirmed using the Plant Phire Direct Kit (ThermoFisher #F-160S) and primers (Supplemental Table 1) as described in LotusBase. Those identified were kept for seed bulking in the previously described growing conditions.

### Bacterial Strains and Culture

*Mesorhizobium loti* R7A and kanamycin-resistant *M. loti* pGingerRFP (red fluorescent protein) were grown at 28°C on tryptone-yeast (TY) media.(22) GV3101 *Agrobacterium tumefaciens* strains pCL2/pCM1/pNOS-mNeonGreen::*LjXLG1/2* were grown at 30°C on Luria Broth (LB) media with 100 µg/mL rifampicin, 30 µg/mL gentamicin, and 50 µg/mL kanamycin.

### RNA extraction and quantitative real-time PCR of target genes

To quantify expression of target genes, 100 mg of roots from the sand cone-tainer rhizobia-inoculated experiment were flash frozen in liquid nitrogen. Total RNA was extracted using the RNeasy Plant Mini Kit (Qiagen) and corresponding DNase. Complementary DNA synthesis was conducted using the SuperScript IV Reverse Transcriptase (Thermo Fisher Scientific) from 500 ng of total RNA, and quantitative polymerase chain reaction (qPCR) was conducted from cDNA diluted 1:5 using the PowerUp SYBR Green Master Mix (Thermo Fisher Scientific). A 200 nM primer concentration and the following protocol were used for qPCR for all targets: 2 min at 50 °C and 2 min at 95 °C, followed by 39 repeats of 15 s at 95 °C, 15 s at 60 °C and 1 min at 72 °C, and ending with 5 s at 95 °C. A melting curve (55–95 °C; at increments of 0.5 °C) was generated to verify the specificity of primer amplification. Three (7dpi) and four (21dpi) biological replicates and three technical replicates of all targets (*LjSXLG1* and *LjSXLG2*) were quantified for gene expression levels relative to the reference gene, a *L. japonicus* polyubiquitin (LotjaGi5g1v0317900), using the ΔΔCT method. All primer sequences used for qPCR can be found in Supplementary Table 1. Raw ΔΔCT values used for statistical analysis can be found in Supplementary Table 2.

### Plasmid Construction and Bacterial Transformation

All expression constructs were generated using PCONS plasmids possessing a GFP-dropout.(23) A monomeric NeonGreen fluorescent protein was fused to the C-terminus of the coding sequences for *LjSXLG1/2* via Gibson reaction (New England Biolabs HiFi Assembly). The fusions were then inserted via Golden Gate BSA1 restriction cloning (New England Biolabs) into two different strength PCONS (pCL2 and pCM1) and a similar plasmid using the nopaline synthase (NOS) promoter. Chemically competent XL1-Blue *E. coli* (QB3 MacroLab, USA) grown at 37°C were used in the construction steps before the final versions were electroporated into GV3101 *Agrobacterium tumefaciens*. α-DHA *E. coli* were used to generate *M. loti* expressing a pGinger RFP construct via conjugation. Briefly, respective bacteria were grown, pelleted, then resuspended and aliquoted as a mix on TY plates at 28°C; after 2-4 days the mixed bacteria were streaked onto TY plates selective for the pGinger plasmid. Successful colonies were cultured, surveyed, and sequenced to determine conjugation success. Plasmids were routinely isolated using the Qiaprep Spin miniprep kit (Qiagen, USA), and all primers were purchased from Integrated DNA Technologies (IDT; Coralville, IA)

### Transient Expression in *Nicotiana benthamiana* and Microscopy

Agroinfiltration protocol was adapted from Sparkes et al.(24) Transformed agrobacterium were grown in LB liquid media with 50 μg/mL kanamycin, 50 μg/mL rifampicin, and 30 μg/mL gentamicin to between optical density (OD) 0.6 and 1 before diluting to 0.05 OD_600_ (SXLG::mNeonGreen fusion) or 0.15 OD_600_ (NLS::mScarlet::NLS) in agroinfiltration buffer (10 mm MgCl_2_, 10 mm MES, pH 5.6). *N. benthamiana* plants were grown and maintained in a temperature-controlled growth room at 25°C and 60% humidity in 16:8 h light:dark cycles with a daytime PPFD of ∼120 μmol m_2_^-1^ s^-1^. *N. benthamiana* were germinated and grown in Sungro Sunshine Mix #4 supplemented with ICL Osmocote 14-14-14 fertilizer at 5 mL/L and agroinfiltrated at 29 days of age. Constructs of interest were infiltrated into the fourth leaf (counting down from the top of the tobacco plant) and harvested 3-4 days post-infiltration. Images were taken using a Zeiss LSM710 laser-scanning confocal microscope at 630x magnification.

### Preparation of *Mesorhizobium loti* for Plant Inoculation

*M. loti* strains were grown for 48 h at 28°C, 180 rpm in TY media. The cultures were then transferred to 15 or 50 mL labeled falcon tubes and pelleted using a swing arm centrifuge at 4000 rcf for 10-15 min. The media was decanted and bacteria were resuspended in DI H_2_O before two rounds of washing. The bacteria were resuspended a final time in DI H_2_0 and adjusted to the desired OD_600._

### Destructive Sampling Sand Cone Experimental Setup

*Lotus japonicus* seedlings were germinated as described and at 7 days old transplanted to 4.75-inch cone-tainers (Ray Leach Stuewe and Sons) filled with 0.3 cm^3^ rockwool at the base, then 27.5 mL medium grain sand (Cemex Lapis Lustre Specialty Sands NO#60), and topped with 10 mL fine play sand (SAKRETE). For uninoculated experiments, plants were watered with a 1/2 MS nutrient solution, covered with cling wrap and a humidifier dome, then placed in the growth chamber. They remained covered for 1 week and were then watered with 1/2 MS once a week for the duration of the experiment. Samples were harvested at desired time points by hand, roots were cleaned with DI water, patted dry with a paper towel, then weighed and images were taken for later measurement in ImageJ.

For inoculated experiments, plants were watered with a 1/2 MS without nitrogen solution, covered with cling wrap and a humidifier dome, then placed in the growth chamber. They remained covered for 1 week before flood inoculation with 3 mL of 0.1 OD_600_ of the desired *M. loti* strain per plant and were then watered once a week with 1/2 MS without nitrogen for the duration of the experiment. Following inoculation, samples were harvested at desired time points by hand, roots were cleaned with DI water and assessed for (im)mature nodule formation while remaining in water to prevent desiccation using an Olympus SZX16 stereoscope fitted with an Olympus SZX2-ILLT base for transmitted/oblique illumination, then patted dry with a paper towel. Finally they were weighed and images were taken for later measurement in ImageJ.

### Non-destructive Sampling Nodule Plate Experimental Setup

Seedlings were transferred ∼4-7 days post germination to 50 mL slanted agar plates made with 1/4 B&D (Broughton and Dilworth) nutrient media without nitrogen and 1.5% w/v plant tissue culture agar (PhytoTech Labs A111). Autoclaved plant germination paper cut to fit was wetted with the same nutrient solution and placed over the agar. Autoclaved metal bars to hold the seedlings and shade roots from light were then placed into the plates. After seedlings were transferred the plates were wrapped 3/4 with parafilm with 1/4 Millipore tape (3M) at the top, placed in a black cardstock cover to shield roots from light, and left at the described growing conditions. One week after transplanting, each plate was inoculated with 500 mL 0.05 OD_600_ *M. loti* RFP, re-wrapped, and returned to the growth chamber. Images were taken every 2 dpi using the Amersham 600 (GE) blue and green light channels for 0.5 seconds each. Additional color photos were taken with a standard camera. Images were analyzed in ImageJ to quantify nodule formation and maturation.

### Data Analysis

All quantitative data analysis and visualization occurred in Jupyter notebook using custom Python scripts using the SciPy, NumPy, Pandas, Matplotlib, and Seaborn libraries. The nonparametric Kruskal Wallis test was used for all nodulation experiment data.

## Results

### Symbiosis-responsive SXLGs share conserved features of the XLG N-terminal domain

Utilizing the peptide BLAST and Expression Atlas features of LotusBase, we identified two homologs of *Lotus japonicus* XLG3 with transcriptional expression profiles associated with the establishment of mycorrhizal and root nodule symbioses, SXLG1 (LotjaGi1g1v0106400) and SXLG2 (LotjaGi6g1v0043000), respectively (Figure 1A). Multiple sequence alignment of SXLG and XLG peptides revealed the conservation to be located in the largely uncharacterized N-terminal domain of conventional XLGs and that SXLGs were approximately one-third their size, the sub-domain of highest conservation contained the putative NLS and the cysteine-rich DUF characteristic of XLGs (Figure 1).(3) This highly conserved DUF has the InterPro annotation “Zinc Finger, RING/FYVE/PHD-type”, sharing the closest resemblance to the Plant HomeoDomain (PHD) motif but lacking the histidine pattern.(25)

**Figure 1.**
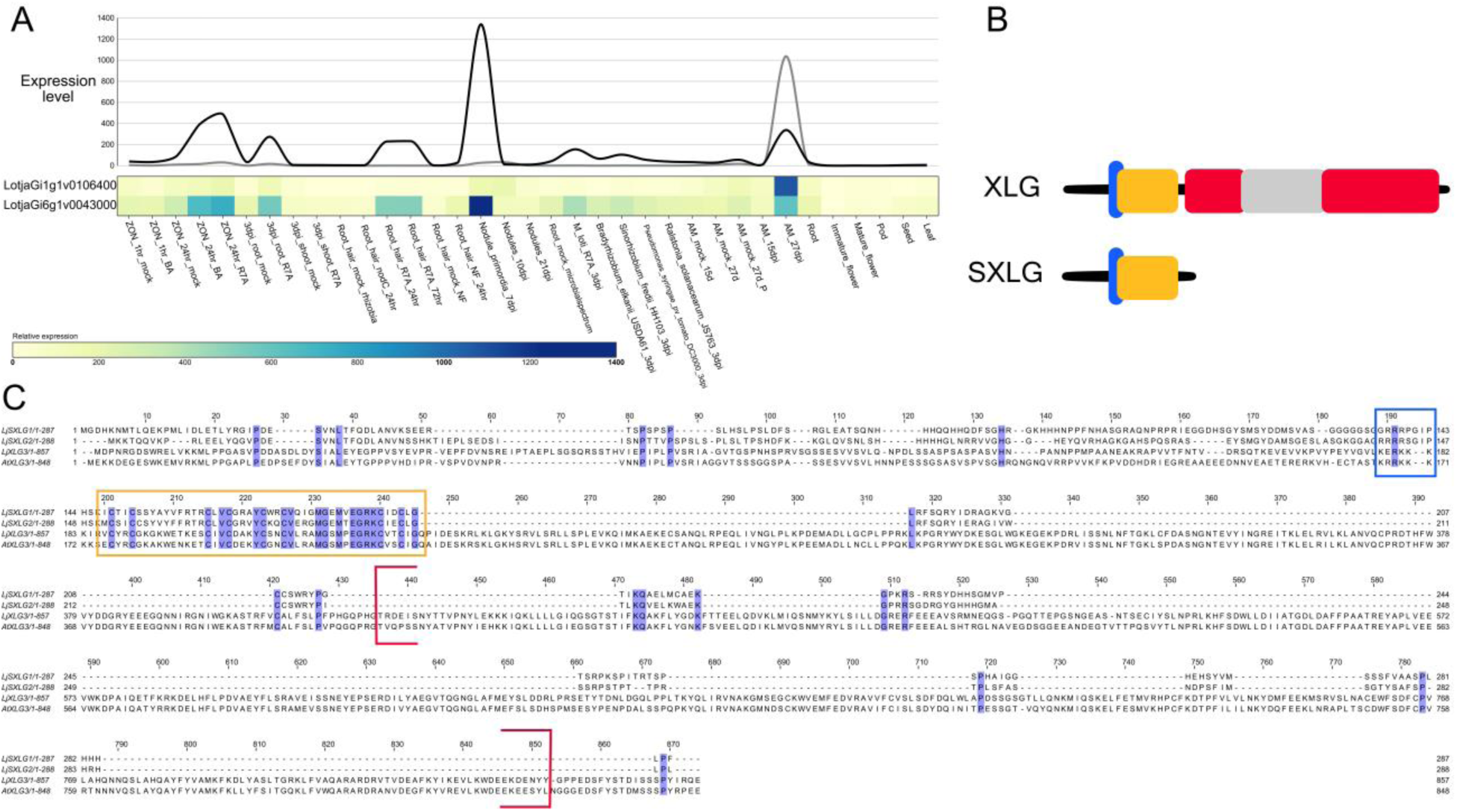
SXLGs share conserved features of the XLG N-terminal domain. **A)** LotusBase Expression Atlas line and heat map displaying the measured expression levels of *LjSXLG1* (LotjaGi1g1v0106400) and *LjSXLG2* (LotjaGi6g1v0043000) across tissues and conditions. Note the tightly controlled patterns: *SXLG1* at 28 dpi mycorrhizal conditions and *SXLG2* also during early stages of the rhizobial symbiosis (recognition and immature nodules) **B)** Graphical representation of XLG domains and the areas SXLGs show the highest conservation, blue = NLS, gold = cysteine-rich domain, red = RAS-like domain, gray = helical domain. Red and gray constitute the Gα-like domain **C)** Muscle peptide sequence alignment of *Lj*SXLGs, *Lj*XLG3, and *At*XLG3. Fully conserved amino acids are highlighted purple, clearly indicating the shared cysteine-rich domain highlighted by the gold box. The NLS is highlighted by the blue box and the Gα-like domain is bracketed in red.

As the size of SXLGs were comparable to the other classes of G-proteins in plants, analysis of InterPro domain annotation in Phytozome and multiple sequence alignments were used to see if they were indeed a class of XLGs. SXLGs did not share significant similarity with the defining domains of other heterotrimeric or small monomeric G-proteins (Supplemental Figure 1).

### Phylogenetic analysis indicates duplication and neofunctionalization of SXLGs in *Fabaceae* species

Many proteins involved with the common symbiotic pathway have undergone duplication and neofunctionalization, with the new copies developing some distinct functions associated with rhizobial symbiosis while older ones retain functions primarily in mycorrhizal symbiosis.(26) As there were two identified SXLGs in *L. japonicus*, each with expression profiles biased towards one symbiosis, it raised the possibility of the genes having undergone this process. Indeed, a phylogenetic analysis indicated such a pattern, with SXLG homologs across *Fabaceae* species forming two distinct clades — one containing the mycorrhizal-responsive SXLG1 and one with the rhizobia-responsive SXLG2 from *L. japonicus* (Figure 2). Additionally, transcriptomic data from MtExpress indicated the *Medicago truncatula* homologs *MtSXLG1* (MtrunA17_Chr5g0432711; Medtr5g075400) and *MtSXLG2* (MtrunA17_Chr3g0107521; Medtr3g467150) grouped according to the same bias in expression profiles.(27) The SXLG1 *Fabaceae* clade was more closely related to putative SXLGs found in plants only able to form mycorrhizal symbiosis (Figure 2).

**Figure 2.**
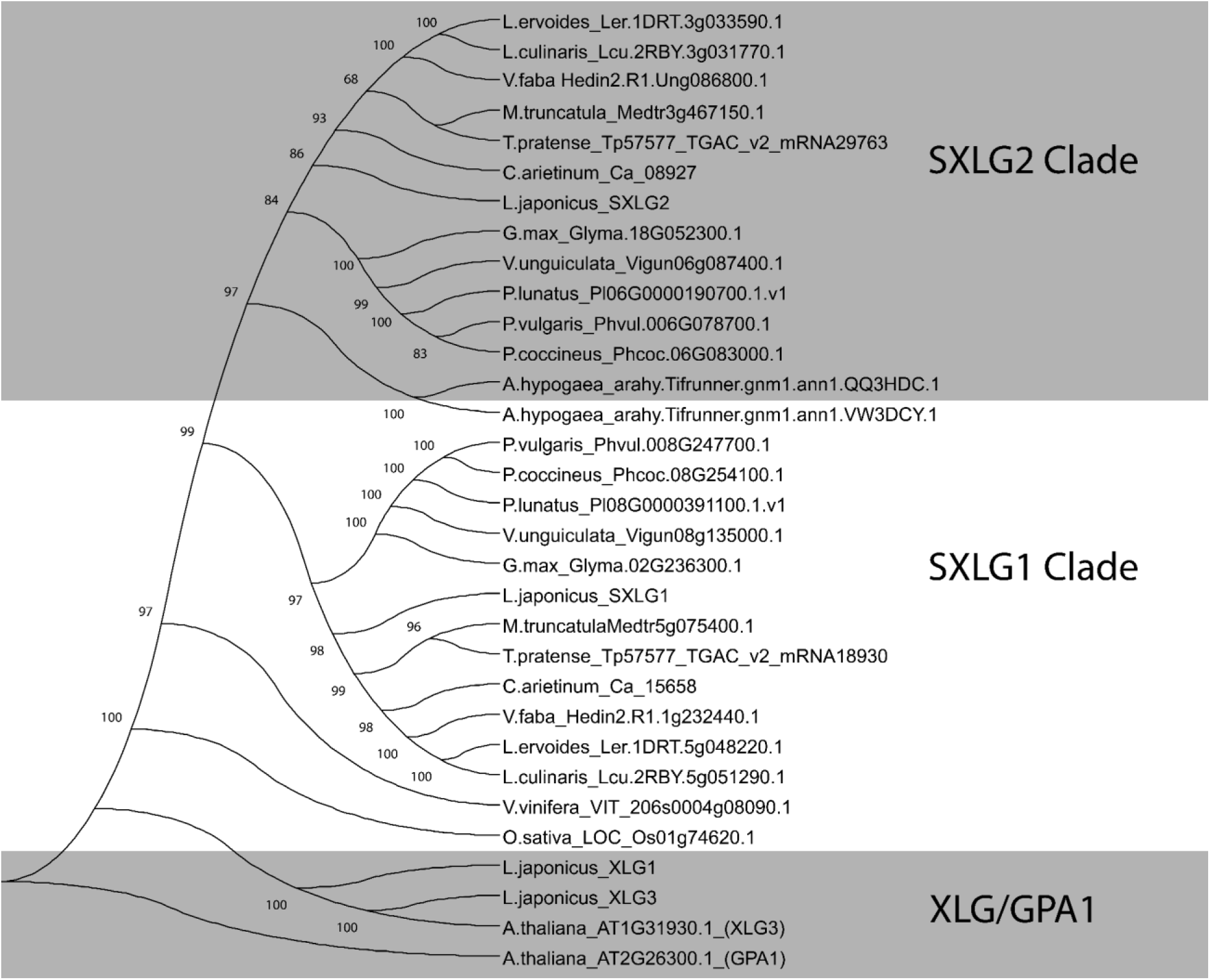
Phylogenetic analysis indicates duplication and neofunctionalization of SXLGs in *Fabaceae* species. SXLGs separate into two clades in *Fabaceae* and away from GPA1 and canonical XLGs. The SXLG1 clade contains non-*Fabaceae* SXLGs and expression data in *L. japonicus* and *M. truncatula* show members to be primarily responsive to mycorrhizal symbiosis. The SXLG2 clade separates furthest from non-*Fabaceae* and expression data show members to be primarily responsive to rhizobial symbiosis. There also seems to be grouping according to nodule type (determinate and indeterminate).

Further supporting this possible duplication event, BLAST results and neighbor-joining tree analysis consistently demonstrated the number of SXLG clades was two in leguminous *Fabaceae* plants and one in plants only able to form mycorrhizal symbiosis (data not shown). Interestingly, *Brassicaceae* species — which are unable to form either symbiosis — were found to lack SXLGs entirely.

### Transient expression reveals cellular localization matching conventional XLGs

XLGs have demonstrated the ability to localize to both the plasma membrane and nucleus, indicating they are not membrane bound like conventional Gα subunits and can traffic through the cytoplasm.(28) Transient expression of *LjSXLG1* and *LjSXLG2* with mNeonGreen fused to the C-terminus in *N. benthamiana* leaves revealed the same capability, with SXLG1 exhibiting reduced nuclear localization compared to SXLG2 (Figure 3). However, both were able to be imported into the nucleolus, something not yet demonstrated in XLGs and absent in our free mNeonGreen control agroinfiltration (Supplemental Figure 3).(3) These results persisted across a range of promoter strengths, infiltration OD_600_, and with mNeonGreen fused to the N-terminus of *LjSXLG*s (Supplemental Figure 3).

**Figure 3.**
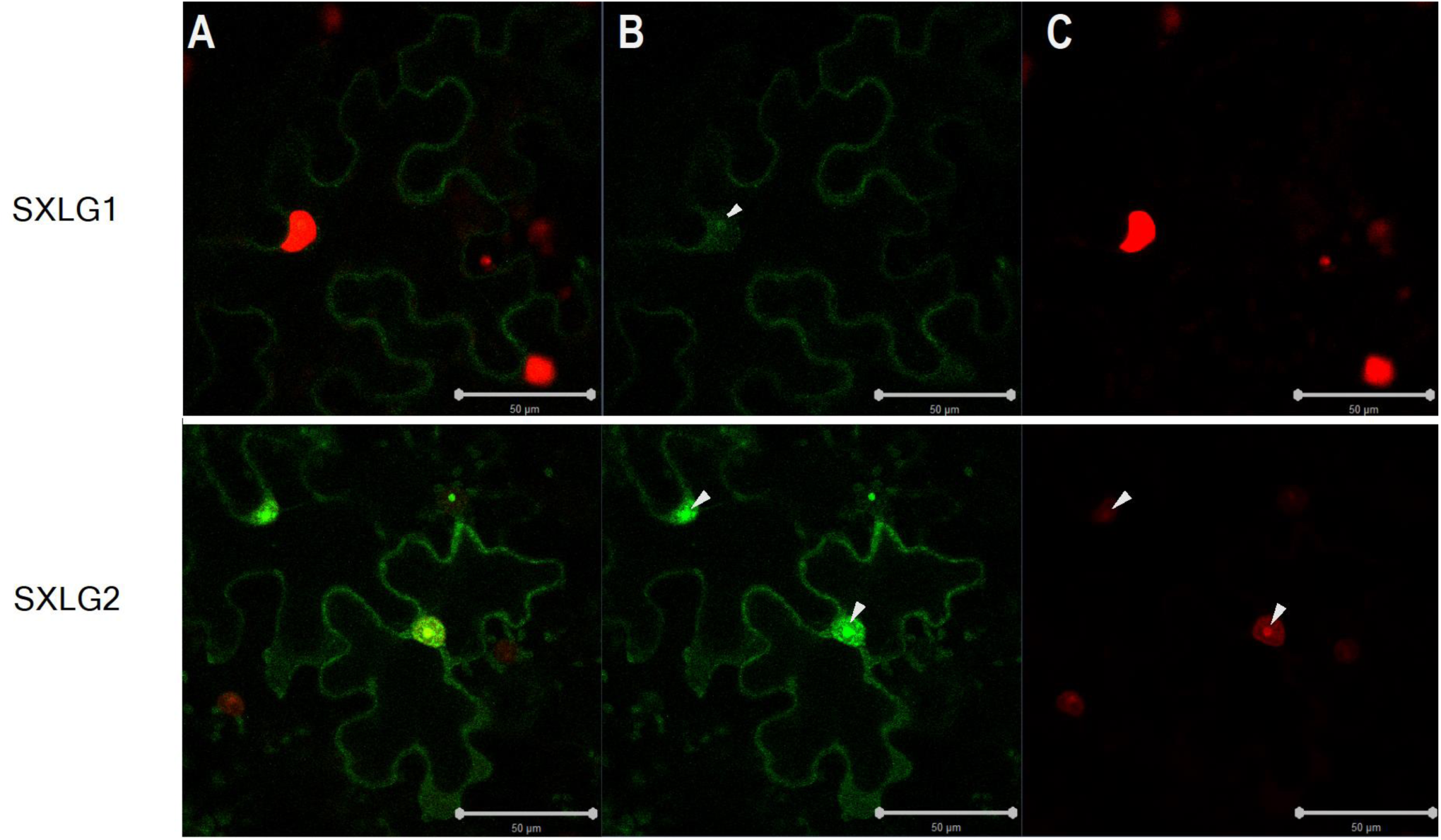
Transient expression reveals cellular localization to plasma membrane, nucleus, and nucleolus. Cells were transfected with a plasmid containing mNeonGreen (mNG) fused to the C-terminus of the coding sequence for *LjSXLG1* (Top) *or LjSXLG2* (Bottom), driven by the pCM2 promoter. An additional plasmid containing mScarlet flanked by a nuclear localization signal is utilized as a nuclear marker. Results were consistent across promoter strength, infiltration OD, and with a N-terminal mNG fusion (Supplemental Figure 3) **A)** Overlay of **B)** LjSXLG::mNeonGreen fusion and **C)** nuclear marker images. LjSXLGs are localized to the plasma membrane, nucleus, and nucleolus. Nucleoli are marked by white arrows. Bars = 50 µm

### Destructive sampling reveals transient impairment of immature nodule formation in *Lotus japonicus sxlg2* mutant

LotusBase ExpressionAtlas shows *LjSXLG2* (LotjaGi6g1v0043000) is primarily expressed in immature nodules and to a lesser extent in early stages of association with rhizobia.(15) Wild type Gifu and transposon knockout mutant plants were grown in sand cones and inoculated with *M. loti* R7A to assess potential effects to nodulation. At 7-and 14-days post inoculation (dpi) *sxlg2* mutants showed fewer immature nodules (*P* ≤ 0.02) and maintained a depressed number, but not statistically significant difference, through the remainder of the experiment compared to Gifu (Figure 4A). However, this only translated into a reduction of mature nodules at 21 dpi (*P* ≤ 0.02) (Figure 4B). There were no observed differences in the size, coloration, or structure of nodules (data not shown).

**Figure 4.**
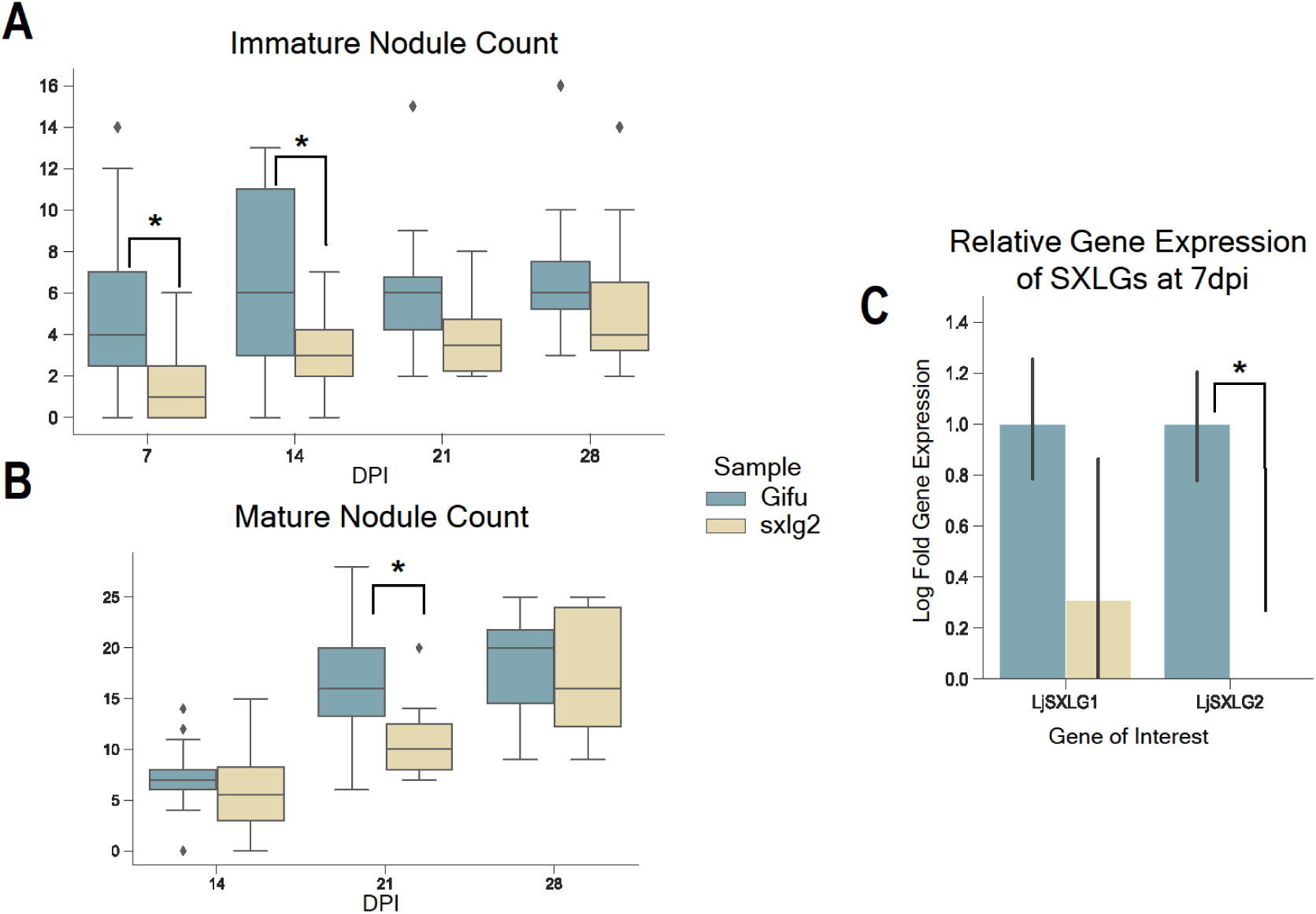
*L. japonicus sxlg2* mutants have transient impairment of immature and mature nodule formation. Comparison of wild type Gifu and *Ljsxlg2* plants grown in nitrogen-depleted conditions in sand cones. **A)** Immature nodule count. n = 19 (7 dpi), 25 (14 dpi), and 10 (21 and 28 dpi) plants. **B)** Mature nodule count. n = 25 (14 dpi), and 10 (21 and 28 dpi) plants. The box plot represents the spread and skewness of data for a given condition by calculating quartiles and determining outliers — the longer the box, the larger the spread of data points from the overall median. The box indicates the Interquartile Range (IQR) by containing the middle 50% of the data, using the medians of the upper and lower halves of the dataset as respective boundaries. The line inside the box indicates the overall median of the dataset. The whiskers (the bars extending from the box) show the range of data within 1.5 times the IQR ending at observed data points termed the maximum and minimum. Individual points shown outside of the whiskers represent outliers within the dataset. **C)** Expression levels of *LjSXLG1* and *LjSXLG2* using *LjUBQ* (LotjaGi5g1v0317900) as reference. n = 3. Error bars indicate 95% confidence intervals. All P-values ≦ 0.02 are denoted by * and calculated by Kruskal-Wallis Test.

### Non-destructive observations of *Lotus japonicus sxlg2* mutants show a reduced number and delay in appearance of established infection events with a potential impact on nodule maturation

Considering the transient nature of the immature nodule phenotype and its inconsistent translation into a reduction of mature nodules, we followed up with non-destructive observations to see if destructive sampling was obscuring important dynamics of the symbiotic infections. Gifu and *sxlg2* mutants were grown on low-nutrient agar plates and inoculated with *M. loti* expressing red fluorescent protein to enable continuous observation. The limited nutrient content also restricted the plants to approximately 2-3 waves of established infection events (where nodules appeared), preventing successive waves from compensating for potential impacts to the early stages of the nodulation process. Images were taken at 2 day increments to measure the number of established infection events and determine the amounts of immature and mature nodules at a given time, based on the visible presence of leghemoglobin causing a pink coloration, with the goal of determining if nodules in *sxlg2* mutants had altered maturation rates or if there was another possible explanation for results seen in destructive sampling experiments. Through 30 dpi *sxlg2* mutants formed less established infection events than Gifu (*P* < 0.02), and when distinguishing mature nodules the reduction was maintained through 40 dpi (Figure 5A, Supplemental Figure 4). This reduction was apparent from the onset of the experiment, where *sxlg2* mutants had a delayed response to inoculation evidenced by less immature nodules through 6 dpi (*P* < 0.02) and less mature nodules from 8 dpi until 40 dpi (*P* < 0.02) (Figure 5B, Supplemental Figure 4). For the duration of the experiment, except for the last two time points (38 and 40 dpi), the *sxlg2* nodules also showed a potential delay in maturation, indicated by a lower percentage of mature nodules (Figure 5C).

**Figure 5.**
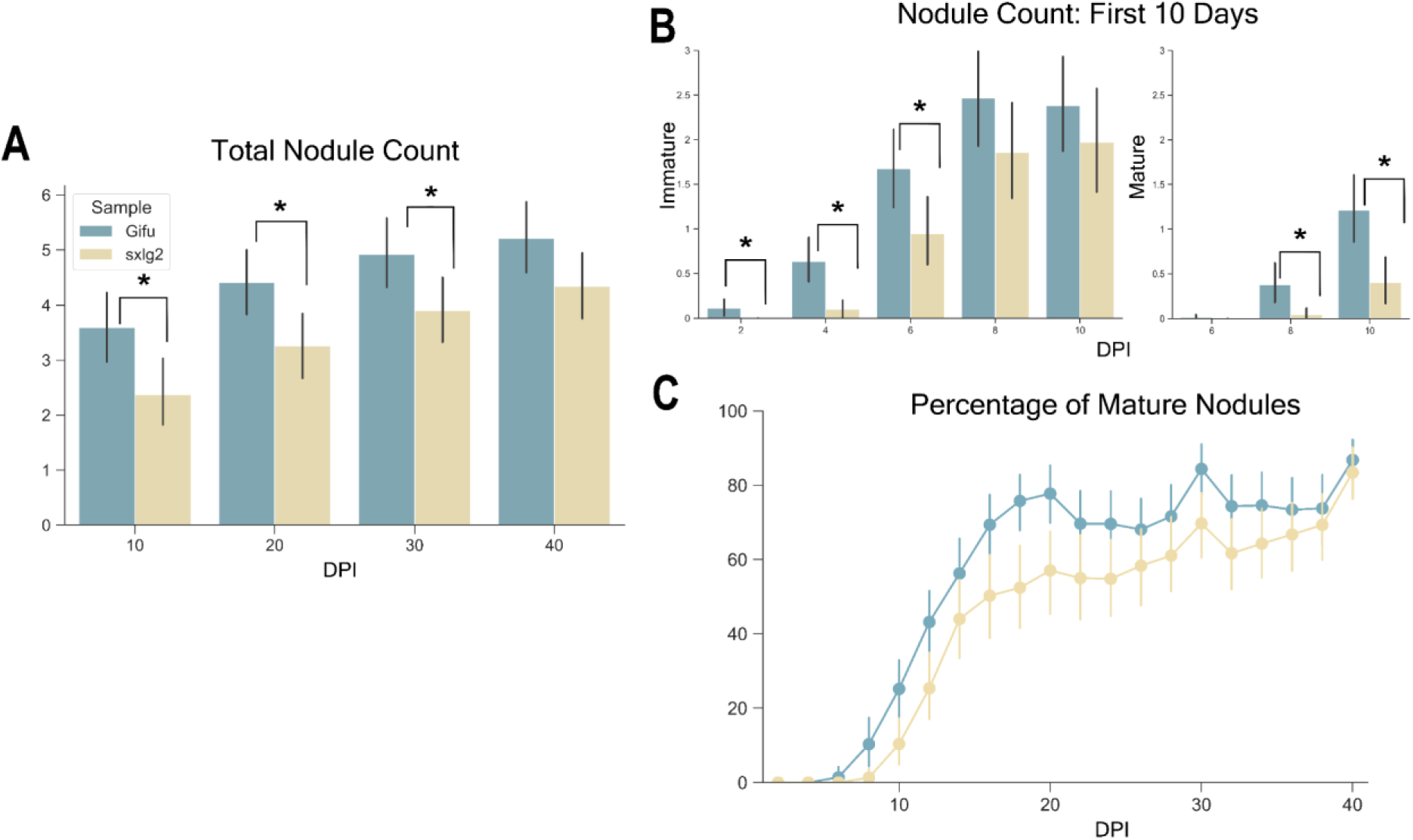
*L. japonicus sxlg2* mutants show a reduced number and delay in appearance of established infection events with a potential effect on nodule maturation. **A**) Comparison of total nodule counts between wild type Gifu and *Ljsxlg2* plants grown in nitrogen-depleted conditions on plates that allowed for non-destructive observation. **B)** The percentage of nodules that were mature at time of sampling. Error bars indicate 95% confidence intervals. n = 70 plants. P-values ≦ 0.02 are denoted by * and calculated by Kruskal-Wallis Test.

### *Lotus japonicus sxlg2* mutants exhibit potential impairment of root growth response to nitrogen limitation

Plants will preferentially allocate resources to grow their root systems when facing nitrogen limitation.(29,30) Physiological root data in both the destructive and non-destructive experimental settings indicate a potential impairment of the exploration response to nitrogen starvation. In the sand cone-tainer nitrogen-restricted experiments primary root length in *sxlg2* mutants did not show the increase at 14 dpi compared to nutrient sufficient conditions that was seen in Gifu (Figure 6B). Additionally, in the nitrogen-restricted non-destructive experiments *sxlg2* mutants had reduced lateral root branching (Figure 6C).

**Figure 6.**
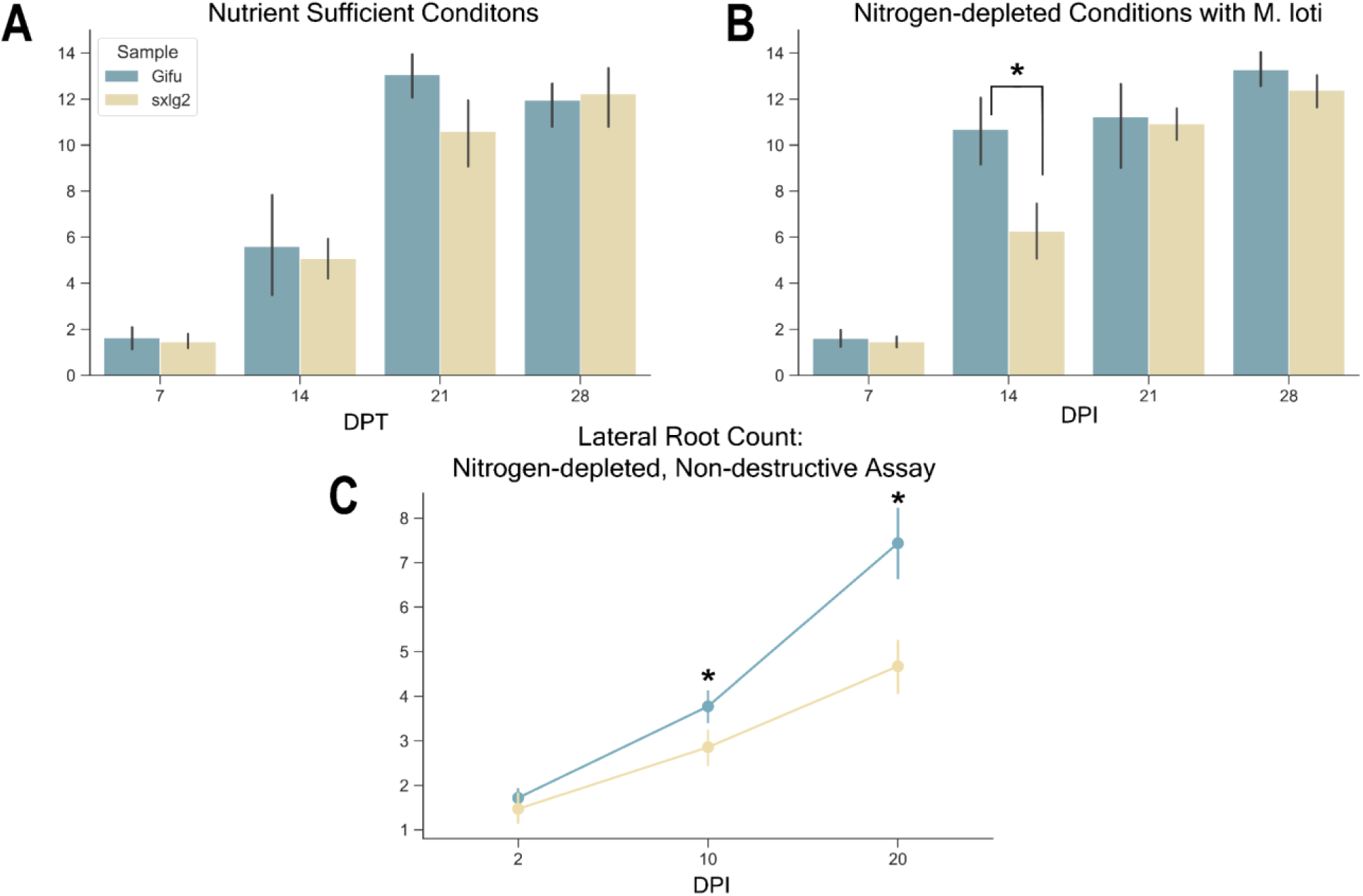
*L. japonicus sxlg2* mutants exhibit potential impairment of root growth response to nitrogen limitation. Comparison of primary root length between wild type Gifu & Lj*sxlg2* plants grown in **A)** Nutrient sufficient, uninoculated conditions (DPT = days post-transplant) and **B)** Nitrogen depleted, *M. loti* inoculated conditions in sand cones. n = 8-10. **C)** Lateral root count through 20 dpi in the nitrogen-depleted, non-destructive conditions on plates inoculated with *M. loti*. Error bars indicate 95% confidence intervals. n = 70 plants. P-value ≦ 0.02 denoted by *. Calculated by Kruskal-Wallis Test.

## Discussion

### SXLGs may provide insight into the evolutionary history and molecular functions of XLGs

SXLGS share the strong conservation of the cysteine-containing DUF seen across XLGs and do not seem to share conserved features found in other classes of plant G-proteins, whether mono– or heterotrimeric, indicating they may be a novel subclass of XLGs (Figure 1, Supplemental Figure 1).(1,2,31) Additionally, similar to XLGs being a class of proteins unique to plants (32), pBLAST results for both *Lj*SXLGs across the UniProtKB/Swiss-Prot and non-redundant protein sequences databases only provided potential homologs among terrestrial plants and some algae. Nearly all of these were either characterized XLGs, annotated as putative XLGs or XLG-like, or uncharacterized proteins. Entering only the cysteine-rich DUF provided the same results, with the exception of three hits; oomycetes and fungi that are either pathogenic or symbiotic to plants. Further, the potential diversity of SXLG homologs (one to two copies per diploid genome) more closely resembles heterotrimeric G-protein subunit diversity (one to four copies per diploid genome) than the typical wide diversity of monomeric G-proteins.(1,33)

Considering that there is currently no known function of this highly conserved cysteine-rich domain and it is believed XLGs arose in plants due to rapid evolutionary steps, such as a potential fusion of the C-terminal Gα-like domain with the uncharacterized N-terminal domain, SXLGs present a missing piece of the XLG puzzle.(34) While the methods used in this study cannot definitively determine whether SXLGs represent the protein that fused to the Gα-like domain or one that arose from separation of the N-and C-terminal domains, it is clear that they contain the previously uncharacterized domain found in all XLGs (Figure 1). Further studies of SXLG molecular structure and function could yield critically lacking information about XLGs.

### SXLGs may be involved in transcriptional regulation

This study demonstrated that SXLGs possess a functional NLS (Figure 3). They also share some alignment with the nuclear export signal in XLGs, however potentially critical hydrophobic residues are lacking and further work is needed to confirm its functionality.(3) Interestingly, both *Lj*SXLGs were shown to localize to the nucleolus in our investigations, which has yet to be demonstrated in XLGs and was absent in our free mNeonGreen control (Figure 3, Supplemental Figure 3). The nucleolus is a site where various regulatory processes occur, such as those involved in the cell cycle and pre-mRNA splicing.(35) The cysteine-rich DUF characteristic of XLGs is annotated as a zinc finger domain and, excluding the histidine residues, shares close resemblance to the Plant HomeoDomain (PHD) motif, which is commonly associated with chromatin modifiers and various transcription factors in plants.(25,36)

The ability to be present at the plasma membrane and in the nucleus and nucleolus, alongside the cysteine-rich domain’s resemblance to the PHD motif, is intriguing evidence of SXLGs potential to contribute to transcriptional regulation in some form. Lending to this hypothesis, XLG2 has been shown to be involved in transcriptional regulation via interaction with RELATED TO VERNALIZATION1 (RTV1) in the nucleus to stimulate chromatin binding.(37) Further study of SXLGs could reveal if they are indeed involved in transcriptional regulation and shed light on the molecular mechanisms other XLGs may be utilizing. This is currently complicated by the lack of a defined protein structure for XLGs beyond the Gα-like domain. Our attempts to generate structures of SXLGs or XLGs *in silico* using AlphaFold resulted in very low-confidence scores outside of a portion of the cysteine-rich domain, which is predicted to form an α-helix (data not shown). Additionally, if SXLGs share the ability to bind to GTP/GDP, Gβγ dimers, or undergo phosphorylation like XLGs is currently unknown. However, XLG2 was shown to be phosphorylated in the N-terminal domain at serine residues upstream the cysteine DUF that *Lj*SXLG1/2 also possess (Figure 1).(3,38)

### SXLGs are likely involved in the establishment of rhizobial and mycorrhizal symbioses

Likely due to their roles in organ development and cell proliferation, many G-proteins have been demonstrated to be involved in various stages of rhizobial and mycorrhizal symbiosis.(4,11) All heterotrimeric G-protein subunits are responsive to or directly involved during symbiosis: Gα were shown to directly interact with NFR1a and NFR1b, Gβ expression levels impact nodule formation, Gγ transcription is up-regulated in both nodulation and mycorrhizal symbiosis, and regulator of G-protein signaling (RGS) expression levels impact nodule formation.(5–7,39) There is also increasing evidence that monomeric small GTPases of the ROP family are influential contributors to infection thread development.(40–42) However, there has been little direct evidence of XLGs involvement in either symbiosis.

A combination of phylogenetic analysis, available transcriptomic data, and experimental observations has indicated that SXLGs may be involved in the establishment of rhizobial and mycorrhizal symbiosis. The potential duplication and neofunctionalization of SXLGs seen in *Fabaceae*, as well as their absence in *Brassicaceae*, indicates involvement in symbiotic relationships (Figure 2). Data from LotusBase and from Frank et al. 2023 have shown *L. japonicus SXLG2* to be expressed during early recognition of compatible rhizobia, in immature nodules, and moved from root hair to cortical cells within the first 10 dpi with rhizobia.(15,43) Similarly, data from multiple studies in MtExpress showed *MtSXLG2* (MtrunA17_Chr3g0107521; Medtr3g467150) to be expressed during nitrogen starvation, early recognition of compatible rhizobia, ectopic addition of Nod factors, and nodule Zones I and II — where infection threads remain present. In Serrano et al. 2024 and MTExpress *MtSXLG1* (MtrunA17_Chr5g0432711;Medtr5g075400) and *MtSXLG2* (MtrunA17_Chr3g0107521;Medtr3g467150) were shown to be expressed in root cortical cells during stages of mycorrhizal association when arbuscules are present.(27,44–50)

These are stages of the symbioses where Nod factor recognition is ongoing and the infection thread (IT) and peri-arbuscular membrane are created and maintained.(51–53) Our experiments demonstrated *L. japonicus sxlg2* mutants to be initially impaired in developing established infection events with the compatible rhizobia *M. loti*, however given sufficient time the *sxlg2* mutants recover from this impairment (Figures 4 and 5). While there may be an impact on the rate at which nodules reach maturity following establishment (Figure 5C), nodules on *sxlg2* mutants were not observed to have differences in size, coloration, or structure. This suggests that SXLG2 is primarily involved in the earliest stages of rhizobial symbiosis, when ITs are formed and migrate into root cortical cells.

This experimental data, the involvement of small GTPases in IT development, mature determinate nodules of *L. japonicus* not maintaining ITs, and *MtSXLG2* expression only in zones of indeterminate nodules where ITs are present, all suggest a possible role of SXLGs in IT development. *SXLG1* expression patterns also indicate potential involvement in peri-arbuscular membrane development or maintenance. However, further work is required to confirm if SXLGs are involved in these processes and the nature of their participation.

## Conclusion

Our study has introduced a novel sub-class of XLGs with conserved features of the N-terminal domain and lacking a Gα-like domain. The cysteine-rich domain within the N-terminus may be involved in transcriptional regulation based on similarity to the PHD domain and SXLG and XLG cellular localization patterns. SXLGs appear to be involved in symbiotic relationships during periods of membrane reorganization; indication of duplication and neofunctionalization in *Fabaceae*, available transcriptomic data in *L. japonicus* and *M. truncatula*, and experiments showing *LjSXLG2* influences the establishment of infection events during rhizobial symbiosis contribute to this hypothesis. SXLGs present an ideal opportunity to better understand the evolution, function, and structure of XLGs and are another example of G-proteins involvement in symbiotic relationships.

## Acknowledgements

Dr. Mitchell Thompson, Dr. Andy Zhou, Dr. Chris Gee, and Liam Kirkpatrick all contributed essential materials and technical support for conducting experiments. Lorenzo Washington was supported by a National Science Foundation Graduate Research Fellowship. This work was supported by the Joint BioEnergy Institute (https://www.jbei.org), U.S. Department of Energy, Office of Science, Biological and Environmental Research Program under Award Number DE-AC02-05CH11231 with Lawrence Berkeley National Laboratory, and by The Novo Nordisk Foundation grant no. NNF19SA0059362 (InRoot).

