## Supplemental Information for "Small Extra-Large GTPase-like proteins influence rhizobial symbiosis in *Lotus japonicus*"

**Supplemental Figure 1. Alignments of SXLGs to other G-protein subunits**

**Supplemental Figure 2. Phylogenetic tree of SXLGs and other G-proteins**

**Supplemental Figure 3. Transient expression of N-terminal fusion of SXLG2 and free mNeonGreen Control**

**Supplemental Figure 4. Non-destructive separated by (im)mature nodules**

**Supplemental Table 1. Primers used in study (qPCR, cloning, genotyping)**

**Supplemental Table 2. qPCR data sheets**

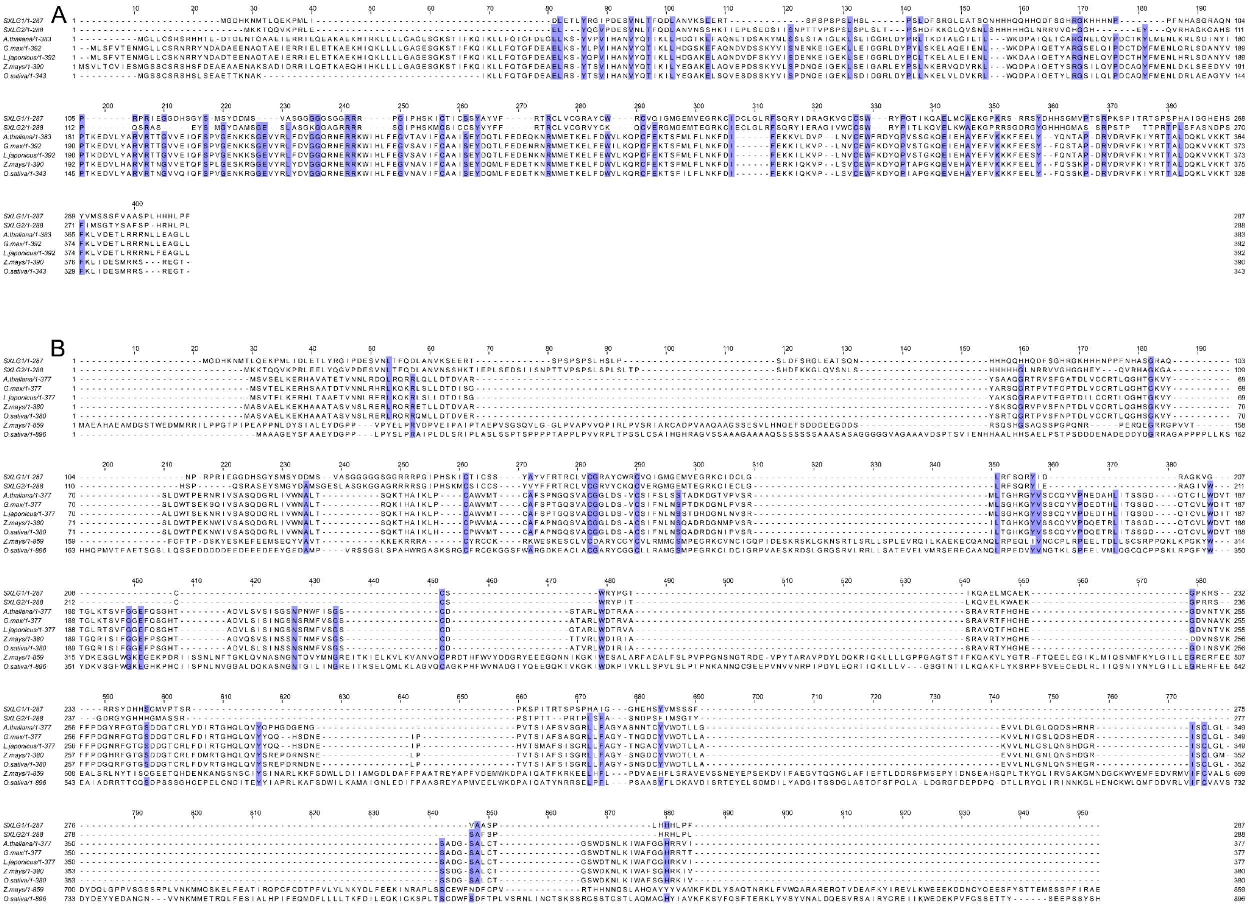

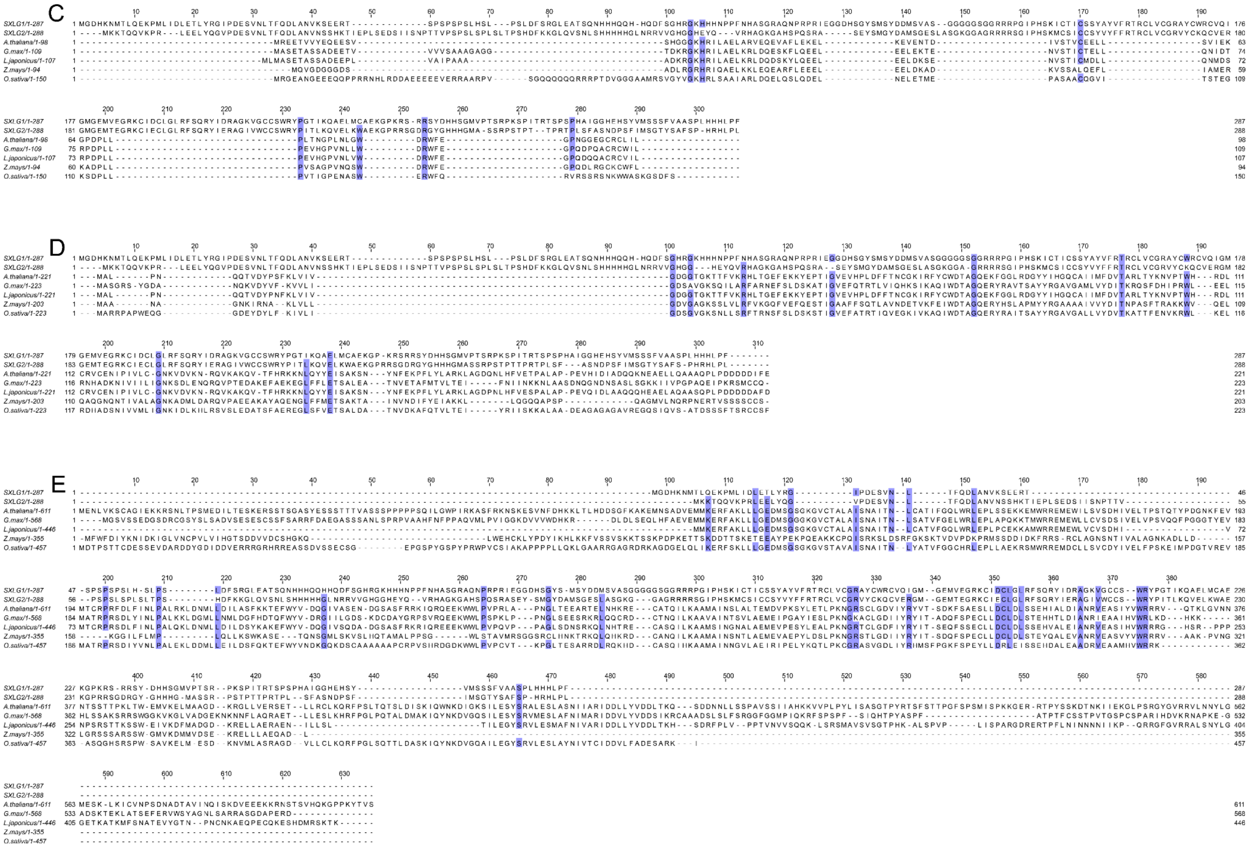


**Supplemental Figure 1. Alignments of SXLGs to other G-protein subunits.** Peptide sequence alignments of LjSXLGs with representatives of **A)** G$\alpha$ subunits **B)** Gβ subunits **C)** G**γ** subunits **D)** Ran GTPases **E)** ROP GTPases from *Arabidopsis thaliana, Glycine max, Lotus japonicus, Zea mays, and Oryza sativa*. The gene IDs are from Phytozome (https://phytozome-next.jgi.doe.gov).

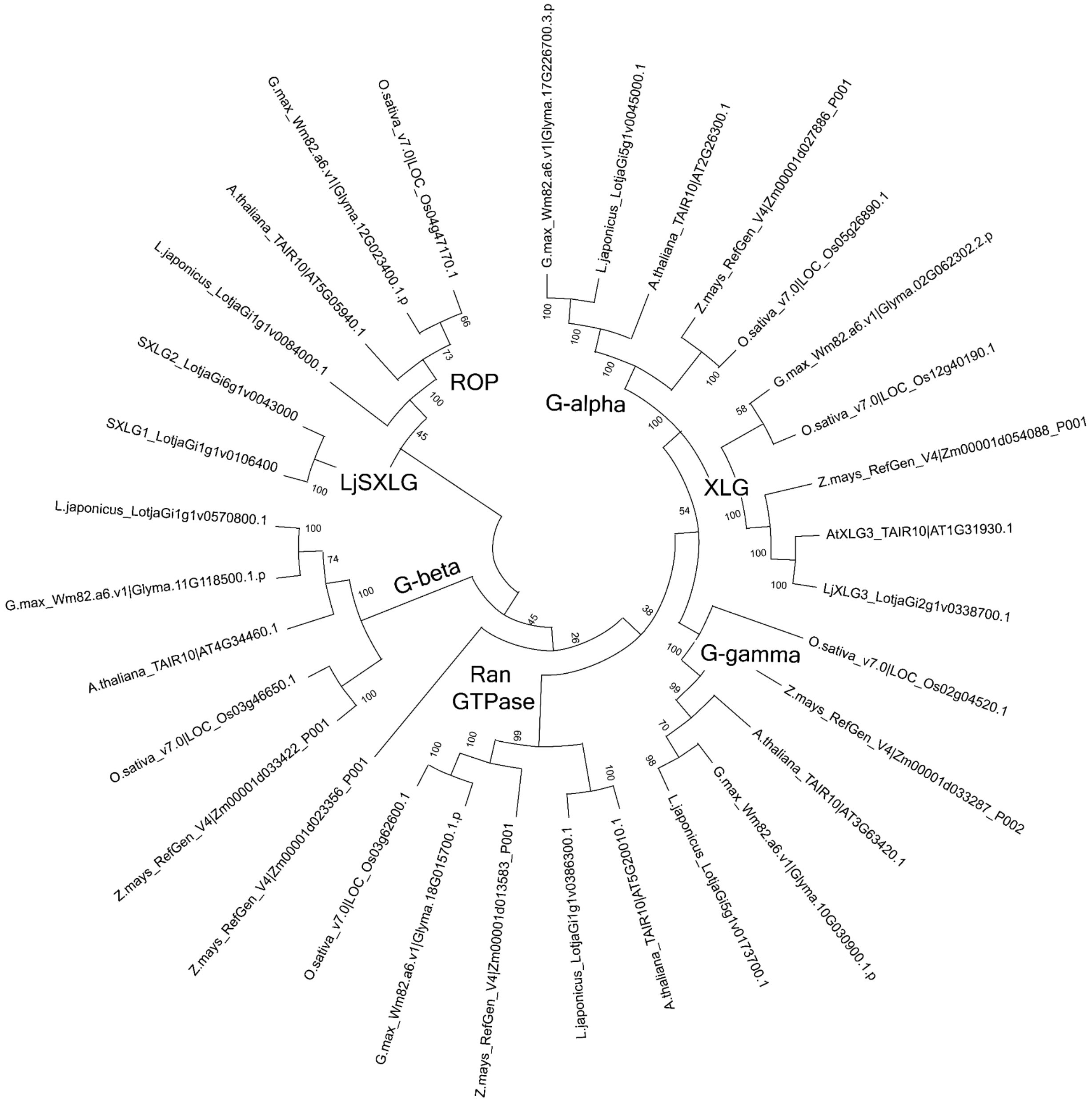


**Supplemental Figure 2. Phylogenetic tree with the G-proteins showing distinct groupings corresponding to Supplemental Figure 1.**

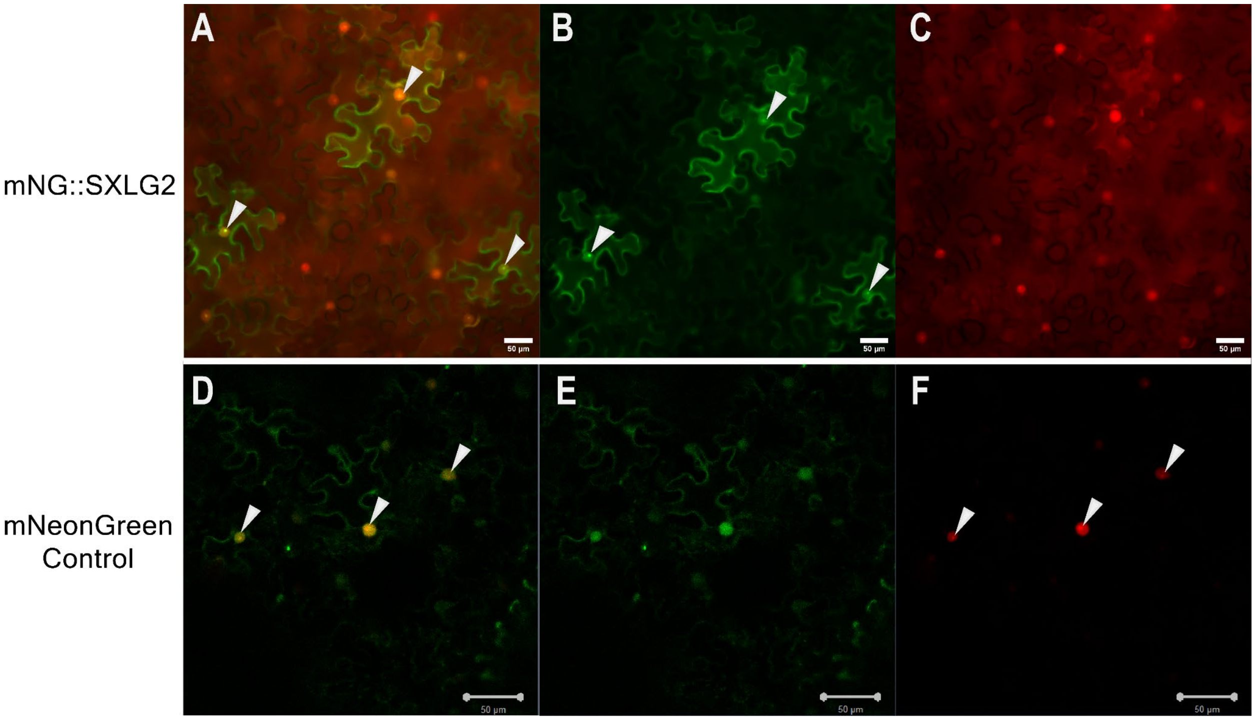


**Supplemental Figure 3. Transient expression of N-terminal fusion of SXLG2 and free mNeonGreen Control.** Cells were transfected with a plasmid containing mNeonGreen (mNG) fused to the N-terminus of the coding sequence for *LjSXLG2*, driven by the pCM2 promoter. An additional plasmid containing mScarlet flanked by a nuclear localization signal was utilized as a nuclear marker. Results were consistent across promoter strength and infiltration OD. **A)** Overlay of **B)** mNeonGreen::SXLG2 fusion and **C)** Nuclear marker. **D)** Overlay of **E)** mNeonGreen expressed without fusion to another protein, driven by pCM2 promoter and **F)** Nuclear marker. Nucleoli are marked by white arrows. Bars = 50 µm

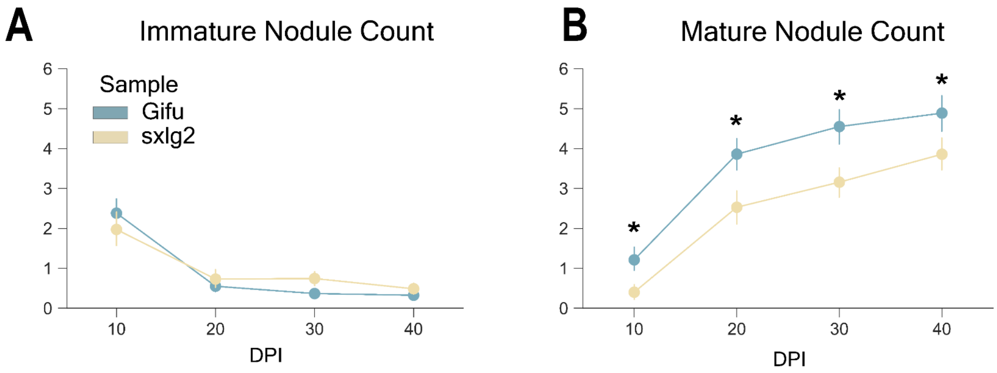


**Supplemental Figure 4. Non-destructive observations separated by (im)mature nodule status.** Comparison of **A)** immature and **B)** mature nodule counts between wild type Gifu & *Ljsxlg2* plants grown in nitrogen-depleted conditions on plates that allowed for non-destructive sample collection. Error bars indicate 95% confidence intervals. n = 70 plants. P-value ≦ 0.02 denoted by *. Calculated by Kruskal-Wallis Test.

#### Supplemental Table 1. Primers used in study

| Treatment | Sample | Gene | log(2^-ΔΔCt) relative to LjUBQ (LotjaGi5g1v0317900) |
|---|---|---|---|
| Gifu | 1_1 | *LjSXLG1* | 0.721451466 |
| Gifu | 1_2 | *LjSXLG1* | 0.500059496 |
| Gifu | 1_3 | *LjSXLG1* | 0.709015077 |
| Gifu | 2_1 | *LjSXLG1* | 1.165545915 |
| Gifu | 2_2 | *LjSXLG1* | 1.100179086 |
| Gifu | 2_3 | *LjSXLG1* | 1.446924002 |
| Gifu | 3_1 | *LjSXLG1* | 1.772923325 |
| Gifu | 3_2 | *LjSXLG1* | 0.774429016 |
| Gifu | 3_3 | *LjSXLG1* | 0.809472616 |
| sxlg2 | 1_1 | *LjSXLG1* | -0.88924662 |
| sxlg2 | 1_2 | *LjSXLG1* | -1.439863943 |
| sxlg2 | 1_3 | *LjSXLG1* | 0.1646334 |
| sxlg2 | 2_1 | *LjSXLG1* | 0.311741992 |
| sxlg2 | 2_2 | *LjSXLG1* | 0.298735974 |
| sxlg2 | 2_3 | *LjSXLG1* | 0.362161927 |
| sxlg2 | 3_1 | *LjSXLG1* | 1.242418752 |
| sxlg2 | 3_2 | *LjSXLG1* | 1.164158951 |
| sxlg2 | 3_3 | *LjSXLG1* | 1.57993786 |
| Gifu | 1_1 | *LjSXLG2* | 0.865146621 |
| Gifu | 1_2 | *LjSXLG2* | 0.430516966 |
| Gifu | 1_3 | *LjSXLG2* | 0.657062642 |
| Gifu | 2_1 | *LjSXLG2* | 1.212531704 |
| Gifu | 2_2 | *LjSXLG2* | 1.103650931 |
| Gifu | 2_3 | *LjSXLG2* | 1.536709442 |
| Gifu | 3_1 | *LjSXLG2* | 1.372680699 |
| Gifu | 3_2 | *LjSXLG2* | 0.845735933 |
| Gifu | 3_3 | *LjSXLG2* | 0.975965062 |
| sxlg2 | 1_1 | *LjSXLG2* | -1.579584099 |
| sxlg2 | 1_2 | *LjSXLG2* | -3.299135425 |
| sxlg2 | 1_3 | *LjSXLG2* | -1.634443531 |
| sxlg2 | 2_1 | *LjSXLG2* | -1.349229901 |
| sxlg2 | 2_2 | *LjSXLG2* | -3.555046122 |
| sxlg2 | 2_3 | *LjSXLG2* | -1.496880123 |
| sxlg2 | 3_1 | *LjSXLG2* | 0.760210442 |
| sxlg2 | 3_2 | *LjSXLG2* | 0.867877382 |
| sxlg2 | 3_3 | *LjSXLG2* | 1.202542896 |

### Supplemental Table 2. Raw log-fold ddCt values for qPCR

| Primer ID | Primer Sequence 5'-3' | Use |
|---|---|---|
| LjUBQ Ref Fwd | GGTCGAAAGCTCTGACACTATT | qPCR |
| LjUBQ Ref Rev | CCTCAAGGGTGATGGTCTTG | qPCR |
| LjSXLG1 qPCR Fwd | GGCCAAAGAGAAGCAGAAGA | qPCR |
| LjSXLG1 qPCR Rev | CTTGTTCGGGTGATAGGACTTT | qPCR |
| LjSXLG2 qPCR Fwd | GGAGGACACGAATATCAGGTTAG | qPCR |
| LjSXLG2 qPCR Rev | TCACCACTCATGGCATCATAG | qPCR |
| P2 | CCATGGCGGTTCCGTGAATCTTAGG | Genotyping |
| sxlg1 Fwd | ACCACCTCCTCCCGAAGCCACACT | Genotyping |
| sxlg1 Rev | TCGACCGGCCTTAGAATTGCCACA | Genotyping |
| sxlg2 Fwd | CCCATGCCTCTTTCCACACACTGC | Genotyping |
| sxlg2 Rev | CACCTTTCAAGACCTTGCCAATGTGAA | Genotyping |
| mUAV Backbone Fwd & Rv | tgccacctgacgtctaagaa | Split destination plasmid to increase amplification for Gibson |
| mUAV-SXLG2 C-term Fwd | TCGGTCTCTAATGAAGAAGACTCAGCAGG | Gibson to make SXLG2::mNG flourescent fusion protein |
| mUAV-SXLG2 C-term Rev | CTGAGTCTTCTTCATAGAGaccgaattcc | "" |
| SXLG2::mNG C-term Rev | CTGACagatccaccacctccAAGAGGGAGGTGACG | "" |
| SXLG2::mNG C-term Fwd | ggaggtggtggatctGTCAGTAAAGGAGAAG | "" |
| SXLG::mNG C-term GG Amp | agggtCTCAAAGCTCAC | Amplify insert to improve efficiency - pairs mUAV-SXLG2 C-term Fwd |
| mNG::SXLG2 N-term Fwd | CAAGggaggtggtggatctATGAAGAAGACTCAGC | Gibson to make mNG::SXLG2 flourescent fusion protein |
| mNG::SXLG2 N-term Rev | agatccaccacctccCTTGTATAACTCATCCA | "" |
| SXLG2-mUAV N-term Fwd | CACCTCCCTCTTTGAGCTTTGAGaccct | "" |
| SXLG-mUAV N-term Rev | gcagggtCTCAAAGCTCAAAGAGGGAGGT | "" |
| mUAV-SXLG1 C-term Fwd | atttctggaattcggtCTCTATGGGTGATCACAAGA | Gibson to make SXLG1::mNG flourescent fusion protein |
| mUAV-SXLG1 C-term Rev | CTTGTGATCACCCATAGAGaccgaattcc | "" |
| SXLG1::mNG C-term Fwd | ggaggtggtggatctGTCAGTAAAGGAGAAG | "" |
| SXLG1::mNG C-term Rev | ctgacagatccaccacctccAAATGGGAGGTGGTG | "" |
| mNG::SXLG1 N-term Fwd | GGGAATGGATGAGTTATACAAGggaggtggtggatctA | Gibson to make mNG::SXLG1 flourescent fusion protein |
| mNG::SXLG1 N-term Rev | GTTCTTGTGATCACCCATagatccaccacctccCTT | "" |
| SXLG1-mUAV N-term Fwd | CTCTCCTCTTCACCACCACCTCCCATTTTAAGCTTTGAG | "" |
| SXLG1-mUAV N-term Rev | gccggactgcagggtCTCAAAGCTTAAAATGGGAGG | "" |
